# Strategic and non-strategic semantic expectations hierarchically modulate neural processing

**DOI:** 10.1101/2020.06.01.127936

**Authors:** Consuelo Vidal-Gran, Rodika Sokoliuk, Howard Bowman, Damian Cruse

## Abstract

Perception is facilitated by a hierarchy of expectations generated from context and prior knowledge. In auditory processing, violations of local (within-trial) expectations elicit a mismatch negativity, while violations of global (across-trial) expectations elicit a later positive component (P300). This result is taken as evidence of prediction errors ascending through the expectation hierarchy. However, in language comprehension, there is no evidence that violations of semantic expectations across local-global levels similarly elicit a sequence of hierarchical error signals – thus drawing into question the putative link between event-related potentials and prediction errors. We investigated the neural basis of such hierarchical expectations of semantics in a word-pair priming paradigm. By manipulating the overall proportion of related or unrelated word-pairs across the task, we created two global contexts that differentially encouraged strategic use of primes. Across two experiments, we replicated behavioural evidence of greater priming in the high validity context, reflecting strategic expectations of upcoming targets based on ‘global’ context. In our pre-registered EEG analyses, we observed a ‘local’ prediction error ERP effect (i.e. semantic priming) approximately 250ms post-target, which, in exploratory analyses, was followed 100ms later by a signal that interacted with the global context. However, the later effect behaved in an apredictive manner - i.e. was most extreme for fulfilled expectations, rather than violations. Our results are consistent with interpretations of early ERPs as reflections of prediction error and later ERPs as processes related to conscious access and in support of task demands.

**Significance statement:** Semantic expectations have been associated with the ERP N400 component, which is modulated by semantic prediction errors across levels of the hierarchy. However, there is no evidence of a two-stage profile that reflects violations of semantic expectations at a single level of the hierarchy, such as the MMN and P3b observed in the local-global paradigm, which are elicited by violations of local and global expectations, respectively. In the present study, we provided evidence of an early ERP effect that reflects violations of local semantic expectations, followed by an apredictive signal that interacted with the global context. Thus, these results support the notion of early ERPs as prediction errors and later ERPs reflecting conscious access and strategic use of context.

## Introduction

Predictive coding theory argues that the brain processes information in a hierarchical probabilistic Bayesian manner (Friston 2005; Knill & Pouget, 2004) by contrasting sensory input with prior expectations generated from context and the perceiver’s knowledge (Heilbron and Chait, 2018; Clark, 2013). Expectations are sent down from higher levels of the hierarchy and any subsequent unexplained sensory input is sent back up the hierarchy as prediction error (Heilbron and Chait, 2018; Friston and Kiebel, 2009; Rao and Ballard, 1999).

Some argue that evoked neural responses (e.g. event-related potentials [ERPs]) reflect prediction errors (Chennu et al., 2013; Friston, 2005). For example, the Mismatch Negativity (MMN)is larger in amplitude for stimuli that do not match short-term auditory expectations, relative to those that do (Heilbron and Chait, 2018). Prediction errors at higher levels of the hierarchy are investigated in paradigms that introduce violations of expectations formed from the global context in which stimuli occur. Indeed, generating such expectations involves complex cognition including working memory and report of conscious expectation (e.g. Bekinschtein et al., 2009). The local-global paradigm (Bekinschtein et al., 2009) elegantly pits local expectation within each trial (i.e. standard vs deviant pitch tones) against a global expectation built from the context across blocks of trials. This paradigm elicits an initial MMN to local violations of expectation, and a subsequent centro-parietal positivity at approximately 300ms post-stimulus (P3b) to global violations of expectation (see Faugeras et al. 2012; King et al., 2013; El Karoui et al., 2015); thereby, separating prediction error signals at two levels of an expectation hierarchy that unfold sequentially.

Within the realm of more ecologically valid stimulus processing, speech comprehension is similarly influenced by expectations at multiple levels of a hierarchy (e.g. Lewis, Bastiaansen, 2015; Ylinen et al., 2016; Lau et al., 2013; Hutchison, 2007; Kuperberg, Jaeger, 2016). The N400 – a negative deflection peaking around 400ms post-stimulus (Kutas, Federmeier, 2011) – is a potential marker of errors of such semantic expectations (Rabovsky & McRae, 2014). On a local level, the N400 is larger to words that have not been primed relative to those that have (e.g. larger for DOG when preceded by Lamp than by Cat; Cruse et al., 2014; Lau et al., 2013; Koivisto & Revonsuo, 2001), and at a more global level, the N400 is larger to words that are unexpected within a sentential context (Brothers et al., 2017; Boudewyn, Long & Swaab, 2015; Thornhill, Van Petten 2012; Van Berkum et al., 1999). Interestingly, unlike the MMN/P3b in auditory processing, semantic prediction errors appear to be reflected in the magnitude of a single component –the N400– rather than in a series of components moving through the hierarchy of relative top-down involvement.

One approach to separate prediction error signals at two levels of a semantic expectation hierarchy is with a prime validity manipulation of a word-pair priming task. Specifically, we can pit the facilitation of target word processing that comes from presentation of a related prime against a global context in which it is not efficient for the comprehender to use the prime to predict the target – i.e. primes rarely followed by related targets (Keefe and Neely, 1990; Hutchison, 2007; Lau et al., 2013(a); Lau et al., 2013(b)). Therefore, as the proportion of related pairs increases within a context, the prime validity increases (i.e. the prime is more likely to predict the target). If individuals use the global context of prime validity to modulate their expectations, behavioural facilitation follows.

In ERP studies of prime validity, this hierarchy of local expectations (i.e. the prime relatedness) and global expectations (i.e. the prime validity) has not been reported to modulate the amplitudes of two sequential components (Boudewyn, Long & Swaab, 2015; Lau et al., 2013); hence, there is no evidence of a two-stage profile to semantic expectation violation. Rather than reflecting error at one level, the N400 (or see Boudewyn, Long & Swaab (2015) for N200 evidence) appears to account for a combination of errors across levels of the hierarchy. To disentangle these results, here we report a pre-registered trial-by-trial manipulation of both local and global semantic expectations.

First, we report a replication of the reaction time facilitation caused by global context as described by Hutchison (2007). Second, we report the associated electrophysiological markers of expectation and violation across levels of the hierarchy from a separate group of healthy participants performing the same task. In accordance with predictive coding, we hypothesised that ERP amplitudes would reflect violations of expectation at consecutive levels of the hierarchy, with local violations evident earlier than global violations.

## Materials and Methods

### Experiment 1 – Behavioural study

#### Participants

We recruited participants through the Research Participation Scheme website of the University of Birmingham, who received credits for their participation. A total of 64 participants were recruited, with the data of two participants excluded from analysis due to outlying data, as quantified by the non-recursive procedure for outlier elimination (detailed below; Van Selst, Jolicoeur, 1994; Hutchison, 2007). Therefore, the final sample consisted of 62 participants (59 female, 3 male; median age: 19, range: 18 – 28). All participants reported to be mono-lingual native English speakers, right-handed, and with no history of neurological conditions or diagnosis of dyslexia. All participants gave written informed consent prior to participation in this study, which was approved by the STEM Ethical Review Committee of the University of Birmingham.

#### Stimuli

Associated prime-target pairs were selected from the Semantic Priming Project database (Hutchison et al., 2013) and the experimental design was a replication of the paradigm implemented by Hutchison (2007). First, all word pairs available in the database (N: 1661) were ordered by Forward Associative Strength (i.e. the proportion of individuals who spontaneously name the same target after reading the prime word) and the 352 word-pairs with the highest strength were selected after removal of any specific American English associations (e.g. Clorox-Bleach; Slacks-Pants).

The first 156 word-pairs from this list of 352 word-pairs with the highest forward association were chosen to be the critical stimuli for statistical analysis. The remaining 196 word-pairs served as fillers to generate the global context and are not included in the statistical analysis. We divided all 156 critical word-pairs into two lists (N: 78 word-pairs per list) that were balanced according to the values from the database (Hutchison et al., 2013) for forward association, length, log HAL frequency, and orthographic neighbourhood (all p>.604; all BF10 <.196). In the same way, we divided the 196 filler word-pairs into two balanced lists (N: 98 word-pairs per list; all p>.284, all BF10<.267). Thus, we had created two critical related word-pair lists and two filler related word-pair lists. To create the unrelated word-pair lists, we manually re-paired (within list) all word-pairs in each of the four lists above (two critical, two fillers) ensuring that unrelated targets were both semantically unrelated to their prime and shared no overlapping phonemes with their respective related target. This resulted in a final set of eight lists: two critical related, two critical unrelated, two filler related, and two filler unrelated. Each participant was assigned two Critical sets of word-pairs (one related and one unrelated; 78 word-pairs per list) and two Filler sets (one related and one unrelated; 98 word-pairs per list). Hence, each participant saw all words within the full set of 352 word-pairs exactly once, composed of 176 related word-pairs and 176 unrelated word-pairs.

To create the prime-validity manipulation, first we assigned half of the Critical word-pairs, including both related and unrelated items, to one colour (yellow or blue), and the other half with the other colour in an interleaved order. Next, the related filler set was assigned with one colour (yellow or blue), and the unrelated filler set was assigned with the other colour. Therefore, across all items seen by each participant, 77.8% of word-pairs presented in one of the two colours were related, thus giving that colour high prime validity, and 77.8% of word-pairs presented in the other colour were unrelated, thus giving that colour low prime validity. Importantly, across the entire set of stimuli that each participant saw, exactly half were related (the other half unrelated) and half were presented in one colour (the other half in the other colour). However, the probability of a related target following a prime of one colour was 77.8% and the probability of a related target following a prime of the other colour was 22.2%. Across participants, the colour assignment of the high validity primes was counterbalanced (i.e. half of participants saw high prime validity word-pairs in blue and low prime validity word-pairs in yellow; and the other half saw the opposite colours for each proportion), and all possible combinations of word lists were used, resulting in 32 permutations.

#### Procedure

The task was presented with Psychtoolbox (Brainard, 1997; Pelli, 1997; Kleiner et al., 2007) in Matlab (Mathworks, Inc., Natick, Massachusetts). The vocal reaction times (RT) were measured with a Cedrus SV-1 Voice Key (Cedrus Corporation), with all participants completing four practice trials under the experimenter’s supervision to adjust the voice key threshold according to the participant’s speech volume. The trial procedure is shown in Figure 1. Specifically, each trial started with a central fixation cross on a grey background lasting 600 ms; then, the prime word was displayed in either yellow or blue, at the centre of the screen for 160 ms; followed by a blank screen for 1080ms, and subsequently the target was displayed on the screen; thus, the stimulus onset asynchrony (SOA) was 1240ms. The target stayed on the screen until the participant pronounced the word; then the word disappeared from the screen, which remained blank for 300ms. Afterwards, a rating for the quality of pronunciation was displayed on the screen with the following questions and potential responses: How would you rate your pronunciation? 1) Correct pronunciation; 2) Unsure of pronunciation; 3) Mispronunciation; 4) Accidental voice-key triggering. Participants gave a button response on the keyboard (1-4) to rate their pronunciation (as per Hutchison, 2007). After the participant responded, the screen remained blank for 1000ms, before the next trial began.

**Figure 1:**
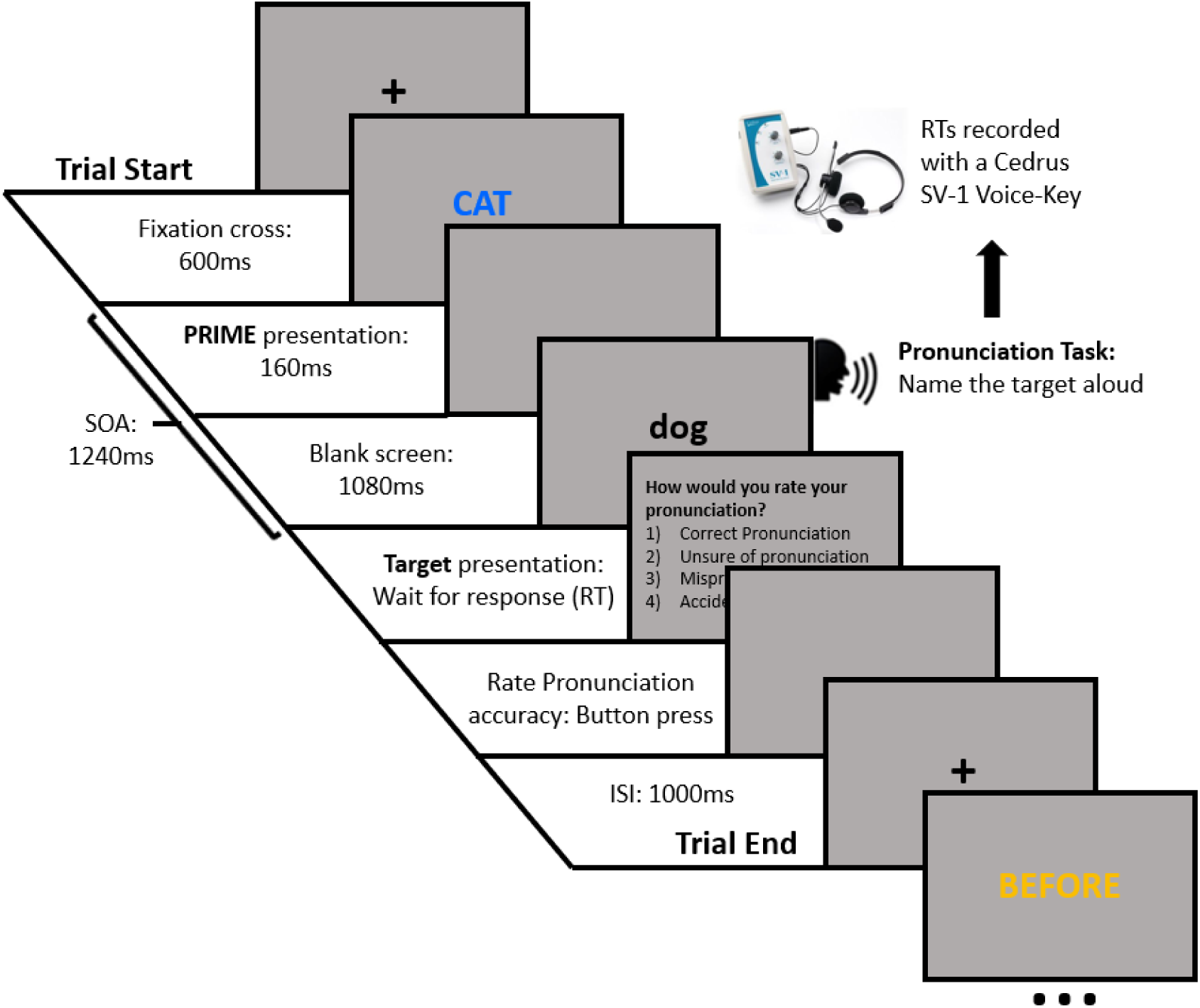
Semantic Priming Relatedness Proportion task (Hutchison, 2007). Participants were required to name the target word aloud and as fast as possible, while their responses were recorded.

Each participant was tested individually and sat approximately 70 cm away from the computer screen. All participants received written information about the study, the instructions and the consent form. In addition, the instructions were verbally repeated by the experimenter. We instructed all participants that a coloured uppercase word (either blue or yellow) will be displayed on the screen and that they must read it silently to themselves; then, a black lowercase word will be displayed on the screen, and they should pronounce the word aloud, as fast and accurately as possible. Participants were told that the colour of the uppercase word will cue the probability of the lowercase target being related or unrelated. Half of the participants received the following written instructions: “If the uppercase word is Blue, it is highly likely that the meaning of the lowercase word will be related; and if the uppercase word is Yellow, it is highly likely that the meaning of the lower-case word will be unrelated” (as per Hutchison, 2007). The other half of participants received the same instructions but with the colours flipped.

After the task, we asked participants to complete a self-report form about the use of strategy throughout the task, to determine whether they were using expectations strategically. The form was composed of three questions and a free text description of the strategy. The questions were the following: 1) Which colour was highly likely to be related? (Responses: BLUE / YELLOW); 2) Did you use the colour of the UPPERCASE word (BLUE, YELLOW) as a cue for knowing whether the following word was related or unrelated? (Responses: YES / NO); 3) Did you engage in any strategy to speed up your responses using the colour cue? (Responses: YES / NO); 4) If YES, briefly describe. We considered participants to have used strategic expectation (i.e. those referred to as the Strategy group) if they correctly identified the colour that was assigned for the high validity condition (Question 1), answered YES in questions 2 and 3, and described a strategy in question 4. All other participants were classified into the No Strategy group.

#### Behavioural Data Analyses

To ensure the inclusion of trials pronounced correctly, we only included trials that were rated by the participants with a correct pronunciation (button press 1); moreover, we eliminated RTs that were longer than 2500ms and shorter than 1ms (i.e. not correctly triggered by the vocal onset). As raw reaction times are skewed, some researchers opt to log transform the data, although this can result in other information about response speed being lost (Lo & Andrews, 2015). Here, we chose to follow the same procedure as in Hutchison (2007) – namely, the non-recursive procedure for outlier elimination (Van Selst & Jolicoeur, 1994). Specifically, reaction times that were more than X standard deviations from the mean were considered to be outliers and were removed, where the value of X decreases with decreasing sample size (i.e. number of trials in each condition for that participant) and is anchored at X=2.5 for a sample size of 100. Next, across all participants we used the same procedure to determine outlier participants and rejected data from two participants that met the outlier criteria. For the remaining 62 participants, a median of 37 trials (range: 16-39) contributed to the high related condition; a median of 36 trials (range: 12-39) to the high unrelated condition; a median of 37 trials (range: 16-39) to the low related condition; and a median of 36 (range: 15-39) contributed to the low unrelated condition.

All behavioural analyses were conducted in Jasp 0.9.1.0 software (JASP Team, 2018). To test for an effect of global context on reaction times, we conducted a two-way repeated measures ANOVA with factors of relatedness (i.e. related vs unrelated targets) and prime validity (i.e. high vs low prime validity). We also reported equivalent Bayesian Repeated Measures ANOVAs (Van Doorn et al. 2019; Wagenmakers et al., 2018). We expected individuals to show faster RTs for related (expected) in contrast with unrelated (unexpected) targets due to local level expectations – i.e. priming. Furthermore, we expected an interaction, with larger priming effects in a high validity context in contrast with a low validity context, reflecting the use of global level context to predict upcoming stimuli.

As a follow-up analysis, we conducted a three-way ANOVA, with its Bayesian equivalent, to test for the interaction and the report of strategy vs no strategy (self-report form) as a between-subjects factor.

### Experiment 2 – Behavioural and electrophysiological study

This study was pre-registered in the Open Science Framework website, details and all codes described in the paper can be found under the following link: https://osf.io/v35te/. Any deviations from the pre-registered methods and analyses are specifically stated in the text.

#### Participants

We recruited participants through the Research Participation Scheme website and placed advertisement posters at the University of Birmingham; participants received a monetary compensation for their participation. We recruited 37 participants, however, since we only investigated those who reported using a strategy, the final sample only included 22 participants (15 female, 7 male; median age: 21, range: 18 - 30; classified by the same report form as experiment 1). The inclusion criteria were the same as those for Experiment 1; however, participants were also required to attend for a structural T1-weighted MRI scan at the University of [name redacted for double-blind review]; therefore, participants who had any metal parts in their body, were claustrophobic, or women who were pregnant were excluded from the study, as the scan was mandatory for participation. All participants gave written informed consent prior to participation in this study, which was approved by the STEM Ethical Review Committee of the University of Birmingham.

We aimed to detect a reaction time interaction of the same magnitude as seen in the Strategy group of Experiment 1; therefore, we conducted a power analysis to select an appropriate sample size for this goal. We performed non-parametric power calculations using the data of all participants of the Strategy group from Experiment 1. Specifically, from the pool of participants of the Strategy group, we selected with replacement N participants and conducted the same two-way repeated measures ANOVA 1000 times to test for the reaction time interaction effect. With an N of 22 participants in the Strategy group we achieved 80% power at p<.05 (i.e. 80% of ANOVAs included a significant interaction).

As we did not know if a participant was in the Strategy group until their self-report form was completed at the end of the study, we recruited participants until 22 of them were classified as being in the Strategy group (median age: 21, range: 18-30; 12 in the no-strategy group, median age: 22, range: 19-33). After removal of trials rated as mispronunciations and those considered outliers according to the non-recursive outlier elimination procedure of Van Selst and Jolicoeur (1994; as Experiment 1), a median of 28 trials (range: 11-38) contributed to the high related condition; a median of 29.5 trials (range: 13-38) to the high unrelated condition; a median of 29 trials (range: 12- 39) to the low related condition; and a median of 28 (range: 14-37) contributed to the low unrelated condition.

#### Stimuli and procedure

Stimuli and procedure were the same as in Experiment 1, except for the duration of the fixation cross (increased from 600ms to 750ms to provide more time for an EEG time-frequency baseline).

#### EEG recording

The EEG signal was continuously recorded with a 125 channel AntNeuro EEG system (Ant-Neuro b.v., Enschede, Netherlands) at a sampling rate of 500 Hz, with impedances kept below 20 kΩ. We placed the ground electrode on the left mastoid bone and referenced online to CPz. As participants were required to pronounce words aloud, we also recorded a bipolar EMG signal with one EMG electrode above the upper lip and the other below the lower lip on the left side of the mouth; approximately over the superior and inferior Orbicularis Oris muscles (Lapatki, Stegeman & Jonas, 2003; Drake, Vogl & Mitchell, 2009).

#### EMG Pre-processing

As this task involved participants speaking, there were considerable artefacts in the EEG data around the vocal reaction time that were challenging to remove adequately. We therefore chose to analyse only the EEG data up to the point of vocal artefact. To minimise artefacts from additional preparatory muscular activity prior to vocal onset, in our pre-registered methods, we planned to choose the latest time-point for analysis post-target by identifying when the mouth EMG signal began to significantly differ between prime validity conditions in a temporal cluster mass randomisation test, as implemented in Fieldtrip (Oostenveld et al., 2011). However, this approach revealed no significant clusters (smallest cluster p = 0.513), and so did not provide a suitable cut-off time-point for our analyses. Therefore, in a deviation from the pre-registered plan, we chose our latest time-point of EEG data to analyse as 150ms prior to the fastest mean RT across conditions (in this instance High Validity – Related = 532ms; see Kuperberg et al., 2018, for a similar approach). Our post-target time-window therefore continued to 382ms post-target. From all the trials included for the statistical analysis only 5.76% of trials had RTs earlier than this time-point, comparable with previous studies (Kuperberg et al., 2018).

#### EEG Pre-Processing Pipeline

We low pass filtered the continuous EEG data at 40Hz using the finite impulse response filter implemented in EEGLAB (Delorme & Makeig, 2004). Due to our interest in analysing slow-waves (see below), we performed no high-pass filtering. Next, we segmented the filtered EEG signals into epochs from 750ms before the onset of the prime up to 382ms post-target (see above for details). Subsequent artefact rejection proceeded in the following steps based on a combination of methods described by Nolan et al. (2010) and Mognon et al. (2011).

First, as in the behavioural data analysis, we excluded all trials in which the participant rated their response as incorrect (i.e. 2, 3, 4 button press) and those that had reaction times that were classified as outliers in the Non Recursive Procedure for outlier elimination (Selst & Jolicoeur, 1994). Next, bad channels were identified and removed from the data. We considered a channel to be bad if its absolute z-score across channels exceeded 3 on any of the following metrics: 1) variance of the EEG signal across all time-points, 2) mean of the correlations between the channel in question and all other channels, and 3) the Hurst exponent of the EEG signal (estimated with the discrete second order derivative from the Matlab function wfbmesti). After removal of bad channels, we identified and removed trials containing non-stationary artefacts. Specifically, we considered a trial to be bad if its absolute z-score across trials exceeded 3 on any of the following metrics: 1) the mean across channels of the voltage range within the trial, 2) the mean across channels of the variance of the voltages within the trial, and 3) the mean across channels of the difference between the mean voltage at that channel in the trial in question and the mean voltage at that channel across all trials. After removal of these individual trials, we conducted an additional check for bad channels, and removed them, by interrogating the average of the channels across all trials (i.e. the ERP, averaged across all conditions). Specifically, we considered a channel to be bad in this step if its absolute z-score across channels exceeds 3 on any of the following metrics: 1) the variance of voltages across time within the ERP, 2) the median gradient of the signal across time within the ERP, and 3) the range of voltages across time within the ERP.

To remove stationary artefacts, such as blinks and eye-movements, the pruned EEG data was subjected to an independent component analysis with the runica function of EEGLAB. The Matlab toolbox ADJUST (Mognon et al., 2011) subsequently identified which components reflect artefacts on the basis of their similarity to stereotypical spatio-temporal patterns associated with blinks, eye-movements, and data discontinuities, and the contribution of these artefact components was then subtracted from the data. Next, we interpolated the data of any previously removed channels via the spherical interpolation method of EEGLAB and re-referenced the data to the average of the whole head.

Before proceeding to group-level analyses, single-subject averages for the ERP analysis were finalised in the following way. First, a robust average was generated for each condition separately, using the default parameters of SPM12. Robust averaging iteratively down-weights outlier values by time-point to improve estimation of the mean across trials. As recommended by SPM12, the resulting ERP was low-pass filtered below 20Hz using a FIR filter (again, with EEGLAB’s pop_newee-gfilt), and the mean of the baseline window (−200 – 0 ms) was subtracted.

Single-subject data for the time-frequency analysis were pre-processed in a similar way. However, first, we concatenated the individual trials into a matrix of channels x all time-points, and filtered each channel in two-steps (high-pass then low-pass) to retain the frequency bands of interest (i.e. 8-12Hz alpha, and 13-30Hz beta), using EEGLAB’s finite impulse response filter (function: pop_eegnewfilt). Next, we extracted the squared envelope of the signal (i.e. the squared complex magnitude of the Hilbert-transformed signal) to provide a time-varying estimate of power within that frequency band. The resulting time-course was re-segmented into its original epochs and averaged within each condition separately using SPM12’s robust averaging procedure. As with the ERP analyses, we low-pass filtered the resulting average time-series below 20Hz (EEGLAB’s pop_newee-gfilt). Finally, we converted the power estimates to decibels relative to the mean of the baseline window (−200 – 0 ms.).

#### EEG / MRI co-registration

We recorded the electrode locations of each participant relative to the surface of the head using a Xensor Electrode Digitizer device and the Visor2 software (AntNeuro b.v., Enschede, Netherlands). Furthermore, on a separate day, we acquired a T1-weighted anatomical scan of the head (nose included) of each participant with a 1mm resolution using a 3T Philips Achieva MRI scanner (32 channel head coil). This T1-weighted anatomical scan was then co-registered with the digitised electrode locations using Fieldtrip.

#### Analyses

##### Behavioural Data Analysis

The behavioural analyses are the same as for the Strategy Group in Experiment 1.

##### EEG Analysis

###### Target ERP, Prime ERP and Prime time frequency analyses

Time-courses (ERPs / time-frequency) within the time-window of interest (0-1240ms for primes; 0-382ms for targets) were compared with the cluster mass method of the open-source Matlab toolbox FieldTrip (Oostenveld et al., 2011). This procedure involves an initial parametric step followed by a non-parametric control of multiple comparisons (Maris, Oostenveld, 2007). Specifically, we conducted two-tailed dependent samples t-tests at each spatio-temporal data-point within our time-window of interest. Spatiotemporally adjacent electrodes (t-values) with p-values < 0.05 were then clustered based on their proximity, with the requirement that a cluster must span more than one time-point and at least 4 neighbouring electrodes, with an electrode’s neighbourhood containing all electrodes within an approximately 4-cm radius (median: 8, range:2-10). Finally, we summed the t-values at each spatio-temporal point within each cluster. Next, we estimated the probability under the null hypothesis of observing cluster sum Ts more extreme than those in the experimental data - i.e. the p-value of each cluster. Specifically, Fieldtrip randomly shuffles the trial labels between conditions, performs the above spatio-temporal clustering procedure, and retains the largest cluster sum T. Consequently, the p-value of each cluster observed in the data is the proportion of the largest clusters observed across 1000 such randomisations that contain larger cluster sum Ts. As our analyses were two-tailed, we set the family-wise error corrected cluster alpha to .025.

###### Prime slow wave linear fit analyses

To further test for ERP evidence of expectation formation in response to the prime, we analysed whether a slow wave differentiates high validity and low validity conditions. For this comparison we used a least-squares linear fit to the averaged ERPs of each condition (High and Low validity primes) for each electrode and participant (as per Chennu et al., 2013). Next, the slope values were compared between conditions with the spatial cluster mass analysis in FieldTrip (Oostenveld et al., 2011).

##### Source estimation analysis

We constructed individual boundary element head models (BEM; four layers) from subject-specific T1-weighted anatomical scans, by using the ‘dipoli’ method of the Matlab toolbox FieldTrip (Oostenveld et al., 2011). Next, we aligned the electrode locations, that were recorded with Xensor Electrode Digitizer device, to the surface of the scalp layer that was segmented from the T1-weighted anatomical scan. For reference points, we used the fiducial points and electrode locations as head shape. We visually checked that the electrode positions and the scalp surface were aligned, and we manually fixed imperfections. We prepared the EEG data before subjecting it to statistical analyses, where we balanced the number of trials in each condition, by taking the smallest condition N as a reference and randomly discarding trials from the other conditions surpassing that N, resulting in equal datasets.

###### ERPs whole brain

For the whole brain ERP source analysis, we used single-trial data that had not been subjected to robust averaging, and defined trials as time windows from −382 to 382ms relative to target onset. This data was then band-pass filtered between 1 and 40Hz using a firws filter as implemented in Fieldtrip (Oostenveld et al., 2011). Subsequently, relative to the different conditions, data were divided into seven sets: one containing all trials, one containing only related trials, one only unrelated trials, one all high-validity related and one all low-validity related trials, one containing all high-validity unrelated and one all low-validity unrelated trials. The sensor covariance matrix was estimated for all these sets of data in the time window −382 – 382ms relative to target onset. A common spatial filter was then computed on the dataset containing all trials using a Linear Constraint Minimum Variance (LCMV) beamformer (VanDrongelen, 1996; VanVeen, 1997; Robinson, 1999). Beamformer parameters were chosen including a fixed dipole orientation, a weighted normalisation (to reduce the center of head bias), as well as a regularisation parameter of 5% to increase the signal to noise ratio (cf. Popov et al., 2018; Sokoliuk et al., 2019). This common spatial filter served then for source estimation of the remaining six sets of trials. Subsequently, the dipole moments of the different source estimates were extracted within the post-stimulus time windows of interest (time windows for source estimates of related vs. unrelated trials: 226-280ms; 232-290ms; 306-382ms; 316-350ms; time window to test interaction effect for source estimates of highly related and unrelated trials and low related and unrelated trials: 316-350ms) and their absolute values averaged over time to obtain one average source estimation value per grid point (VE) and condition.

To test for significant differences between conditions we conducted five contrasts as mentioned above; first, an interaction between prime validity (High/Low) and relatedness of the target (Related/Unrelated) in a time-window from 316 to 350ms; next, we tested the early and late main effects of relatedness of the target (Related/Unrelated) as observed in the sensor analyses results (four main effects), in their respective time windows for the early effect (226-280ms and 232-290ms); and the late effect (306-382ms and 316-350ms). Montecarlo Cluster-based permutation tests were computed as implemented in Fieldtrip (Oostenveld et al., 2011) by using averaged data over each time-window; moreover, we used an alpha and a cluster alpha level of 0.025 and 1000 permutations.

###### Automated Anatomical Labelling (AAL) analysis

We tested for the post-target interaction, between the relatedness of the target (related/unrelated) and the validity of the prime (High prime validity/Low prime validity) in five specific anatomical regions of interest that are defined using the automated anatomical labelling (AAL) atlas (see Brookes et al., 2016; Sokoliuk et al., 2019 for similar analyses with MEG and EEG data). The selected regions are the Left inferior frontal gyrus (LIFG), including pars opercularis, pars triangularis and pars orbitralis; the posterior Left middle temporal gyrus (LMTG); and posterior Left superior temporal gyrus (LSTG), as Weber et al. (2016) reported a relatedness proportion interaction in these regions. In addition, we tested the post-target interaction in the anterior LMTG and anterior LSTG, as Lau et al. (2014) found differences in the anterior left superior temporal region (LSTG) in related vs. unrelated items in a high validity condition. Moreover, as a deviation from our preregistered analyses, we tested the main effects found in the Related – Unrelated contrast at the sensor level (ERPs) in the same anatomical regions (more details in results section). To determine both the anterior and posterior parts of the LMTG and LSTG, we calculated the centre of mass of each AAL region and selected all virtual electrodes that were anterior or posterior to the centre of mass.

We aggregated the AAL regions of interest to each participant’s T1-weighted image; next, for each participant individually, we extracted the average source estimation values of all VEs (from prior source estimation (cf. *ERPs whole brain*)) within each AAL region, weighted them according to their Euclidian distance to the centre of mass of the AAL region (Brookes et al., 2016) and averaged over VEs within each AAL region of interest. We then conducted paired-sample t-tests between the post-target conditions (SP-High validity / SP-Low validity) for all AAL regions; and another paired-sample t-test between the relatedness conditions (Related / Unrelated) for each AAL region in four time windows (226-280ms; 232-290ms; 316-350ms; 306-382ms) from the main effects obtained in the sensor level ERP analyses (results section). The p-values that we obtained were corrected for multiple comparisons across AAL regions using False Discovery Rate, FDR (Yekutieli, Benjamini, 1999). Furthermore, to test for evidence for the null hypothesis, we calculated Bayes Factors using the Bayes equivalent t-test, according to Rouder et al. (2009).

## Results

### Experiment 1 – Behavioural only

In a two-way repeated measures ANOVA, we found a significant interaction between prime validity and relatedness of the target (F (1, 61) = 13.751, p < 0.001, ηp^2^ = 0.184), which was also supported by a Bayesian Repeated Measures ANOVA (BF_inclusion_ = 19.25). As shown in Table 1, this interaction stems from the larger semantic priming effect in the high prime validity context (t (61) = −6.525, p < 0.001, Cohen’s d = −0.829, CI = −1.115 −0.537) relative to the low prime validity context (t (61) = −5.169, p < 0.001, Cohen’s d = −0.656, CI = −0.929 −0.380). Furthermore, reaction times to unrelated items were markedly similar across contexts (t (61) < .001, p = 0.999, Cohen’s d < 0.001, CI = −0.249 0.249), while the difference in semantic priming stems from significantly different reaction times to related items (t (61) = −3.797, p < 0.001, Cohen’s d = −0.482, CI = −0.744 - 0.217).

**Table 1:**
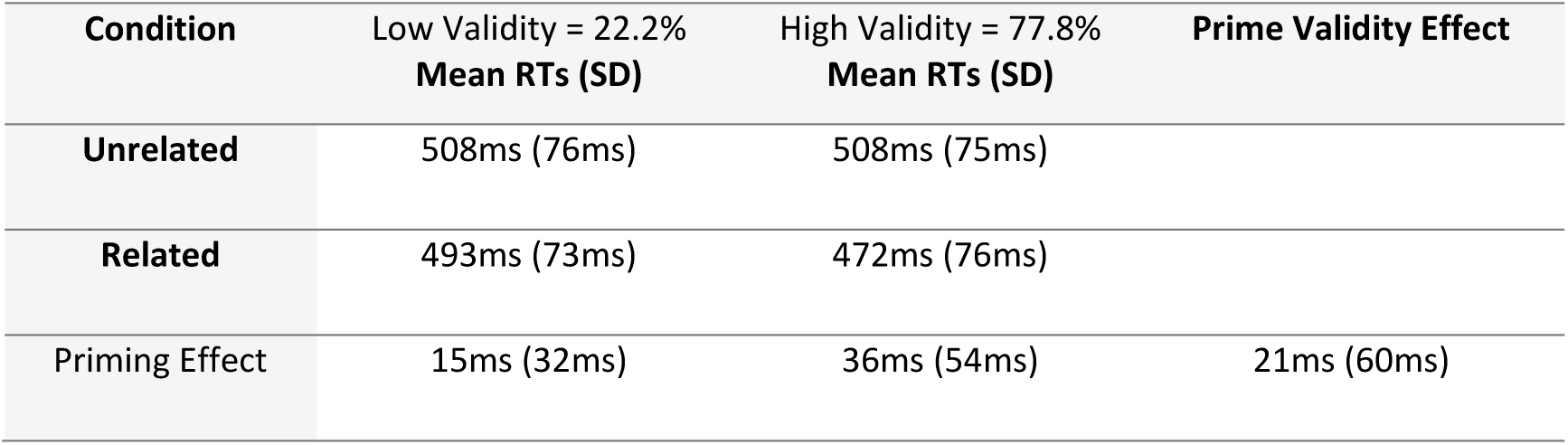
Descriptive statistics including Mean RT (ms) and standard deviation of related and unrelated word-pairs on each validity context, High Prime Validity and Low Prime Validity. Semantic priming effects and prime validity effect (relatedness proportion effect).

| Condition | Low Validity = 22.2%<br>Mean RTs (SD) | High Validity = 77.8%<br>Mean RTs (SD) | Prime Validity Effect |
| --- | --- | --- | --- |
| Unrelated | 508ms (76ms) | 508ms (75ms) |  |
| Related | 493ms (73ms) | 472ms (76ms) |  |
| Priming Effect | 15ms (32ms) | 36ms (54ms) | 21ms (60ms) |

Of 62 participants, 32 were classified in the “No-Strategy” group and 30 were classified in the “Strategy” group. A post-hoc mixed design ANOVA with two within factors (Relatedness of Target; Validity of the prime) and one between subjects factor (Strategy; No-strategy) revealed a significant Target * Prime Validity * Strategy interaction (F (1, 60) = 7.537, p =0.008, ηp^2^ = 0.112, BF_inclusion_ = 3.203), reflecting the apparent presence of a prime validity effect when participants reported using the prime strategically (F (1, 29) = 20.388, p < 0.001, ηp^2^ = 0.413; BF_inclusion_ = 34.67) but absence of a prime validity effect when participants reported no strategy (F (1, 31) = 0.860, p = 0.361, ηp^2^ = 0.027; BF_inclusion_ = 0.393; Figure 2). The No strategy group, however, did exhibit a significant semantic priming effect by showing faster responses in the related relative to unrelated items (F (1, 31) = 21.656, p < 0.001, ηp^2^ = 0.411; inclusion BF_inclusion_ = 4994.57).

**Figure 2:**
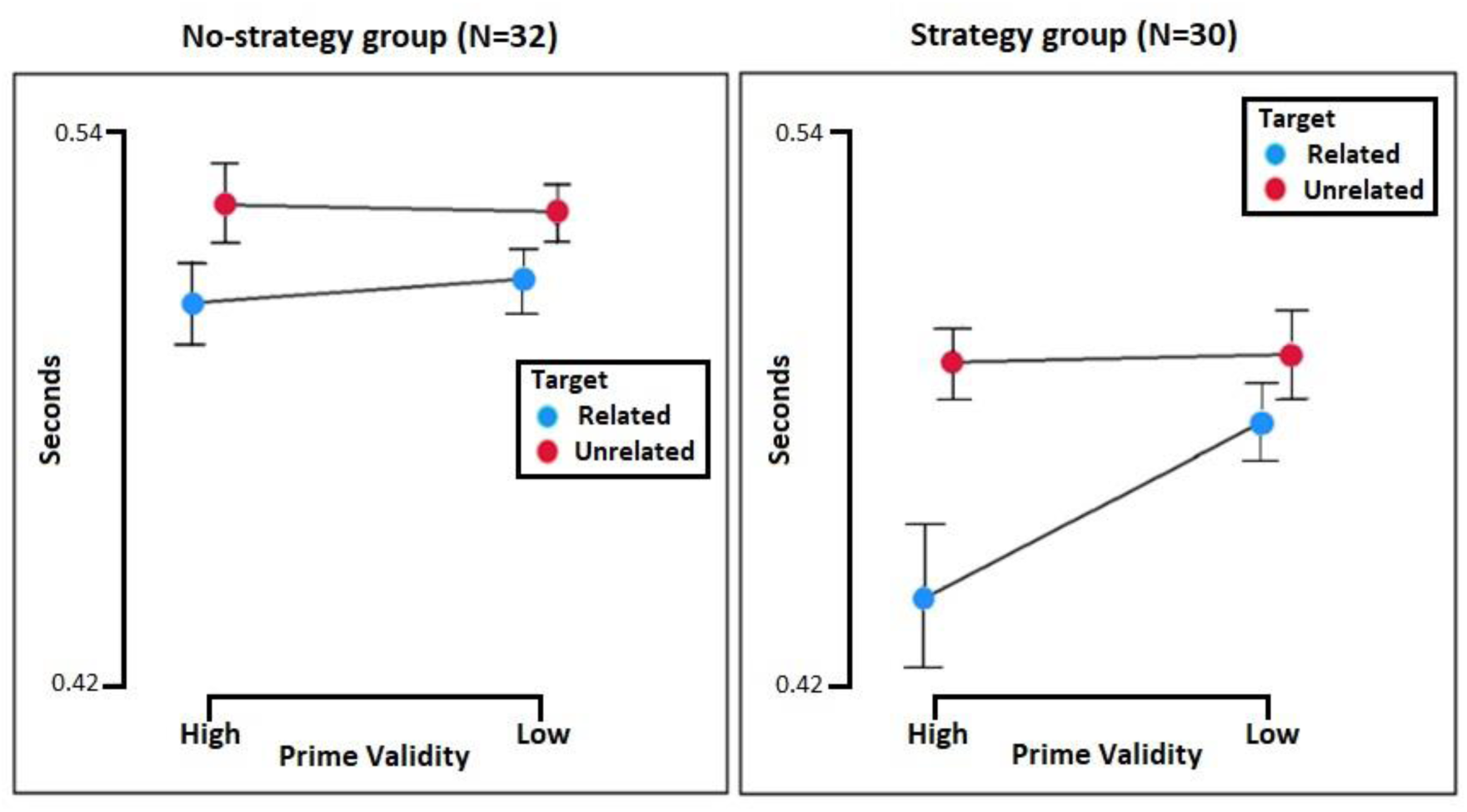
Mean RTs: Prime Validity (High / Low), Relatedness of the target (Related / Unrelated). Interaction (p < 0.001) between the validity of the prime and the relatedness of the target in the group of participants that reported the use of a conscious strategy (right), and no interaction (p = 0.361) in the group of participants that did not report a conscious strategy (left).

### Experiment 2

#### Behavioural Results

These results were qualitatively consistent with those we observed in Experiment 1. A two-way repeated measures ANOVA analysis showed a significant interaction between prime validity and relatedness of the target (F(1, 21) = 9.071, p = 0.007, ηp^2^ = 0.302), while the Bayesian Repeated Measures ANOVA analysis showed anecdotal evidence for the interaction (BF_inclusion_ = 2.519). The interaction was driven by a larger semantic priming effect in the high prime validity context (t (21) = −4.254, p < 0.001, Cohen’s d = −0.907, CI = −1.398 −0.400) than in the low prime validity context (t (21) = −2.046, p = 0.054, Cohen’s d = −0.436, CI = −0.869 0.007), see table 2. There was no significant difference between the reaction times to unrelated items across contexts (t (21) = 0.731, p = 0.473, Cohen’s d = 0.156, CI = −0.266 0.575) as opposed to a significant difference between related items across contexts (t (21) = −2.719, p = 0.013, Cohen’s d = −0.580, CI = −1.027 −0.121).

**Table 2:** Descriptive statistics including Mean RT (ms) and standard deviation of related and unrelated word pairs on each validity context, High Prime Validity and Low Prime Validity. Semantic priming effects and prime validity effect (relatedness proportion effect).

| Condition | Low Validity = 22.2%<br>Mean RTs (SD) | High Validity = 77.8%<br>Mean RTs (SD) | Prime Validity Effect |
| --- | --- | --- | --- |
| Unrelated | 576ms (92ms) | 582ms (87ms) |  |
| Related | 560ms (107ms) | 532ms (110ms) |  |
| Priming Effect | 16ms (54ms) | 50ms (69ms) | 34ms (95ms) |

#### EEG Results – Sensor Level

##### Prime analyses: ERPs, time frequency and slow wave linear fit analyses

As the global context was instantiated by the prime words, we sought to also investigate potential electrophysiological markers of expectation setting (rather than post-target prediction errors). However, none of our pre-registered analyses in the prime time-window (0-1240ms after prime onset) revealed evidence of markers of expectation in response to the prime. Specifically, there were no effects in analysis of the ERPs (smallest cluster p = 0.233), the slow wave linear fit analysis (no clusters formed), or the alpha-beta time-frequency analysis (smallest cluster p = 0.136).

Therefore, in exploratory analyses, we focused the time-window of interest for the ERP analysis on the peak of the global field power (530-1240ms), however this also revealed no significant difference between the high and low validity contexts (smallest cluster p = 0.139). Similarly, we used the window of interest for the alpha-beta time-frequency analysis to the peak of the global field power (602-1240ms), which also yielded no significant difference between conditions (no clusters formed). Moreover, as alpha-beta frequency bands include a wide range of frequencies we analysed them separately. However, the time-frequency analysis in the Alpha band (8-12Hz) showed no significant differences between conditions in the 0-1240ms time window (smallest cluster p = 0.121), nor in the 530-1240ms time window (smallest cluster p = 0.08). The same was true for the Beta band (13-30Hz; 0-1240ms cluster p = 0.312; 530-1240ms cluster p = 0.197). Together, these analyses suggested no apparent electrophysiological markers of pre-target expectation formation in our data.

##### Target Results: ERPs

In our pre-registered interaction contrast in the latency range from 0 to 382ms post-stimulus, the cluster-based permutation analysis yielded no clusters. However, in pre-registered analyses of main effects in the same latency range, we found four significant main effects of relatedness of the target (i.e. unrelated versus related targets; see Figure 3). The clusters in our data occurred in two distinct periods within the time window as shown in Figure 3. Specifically, two clusters reflected a left fronto-temporal dipolar effect of relatedness (Panels A & B in Figure 3) at approximately 250ms post-stimulus (negative cluster: 226 – 280ms, p = 0.019; positive cluster: 232 – 290ms, p = 0.009), and two clusters reflected a later parieto-occipital dipolar effect of relatedness (Panels C & D in Figure 3) at approximately 350ms post-stimulus (negative cluster: 316 – 350ms, p = 0.021; positive cluster: 306 – 382ms, p = 0.004). The early effects showed a predictive signal as in both clusters the voltage exhibited more extreme values for unrelated than related items. On the contrary, the later effects showed signs of an apredictive signal, especially in Panel D, as the voltage within the cluster had more extreme values for the related relative to the unrelated items.

**Figure 3:**
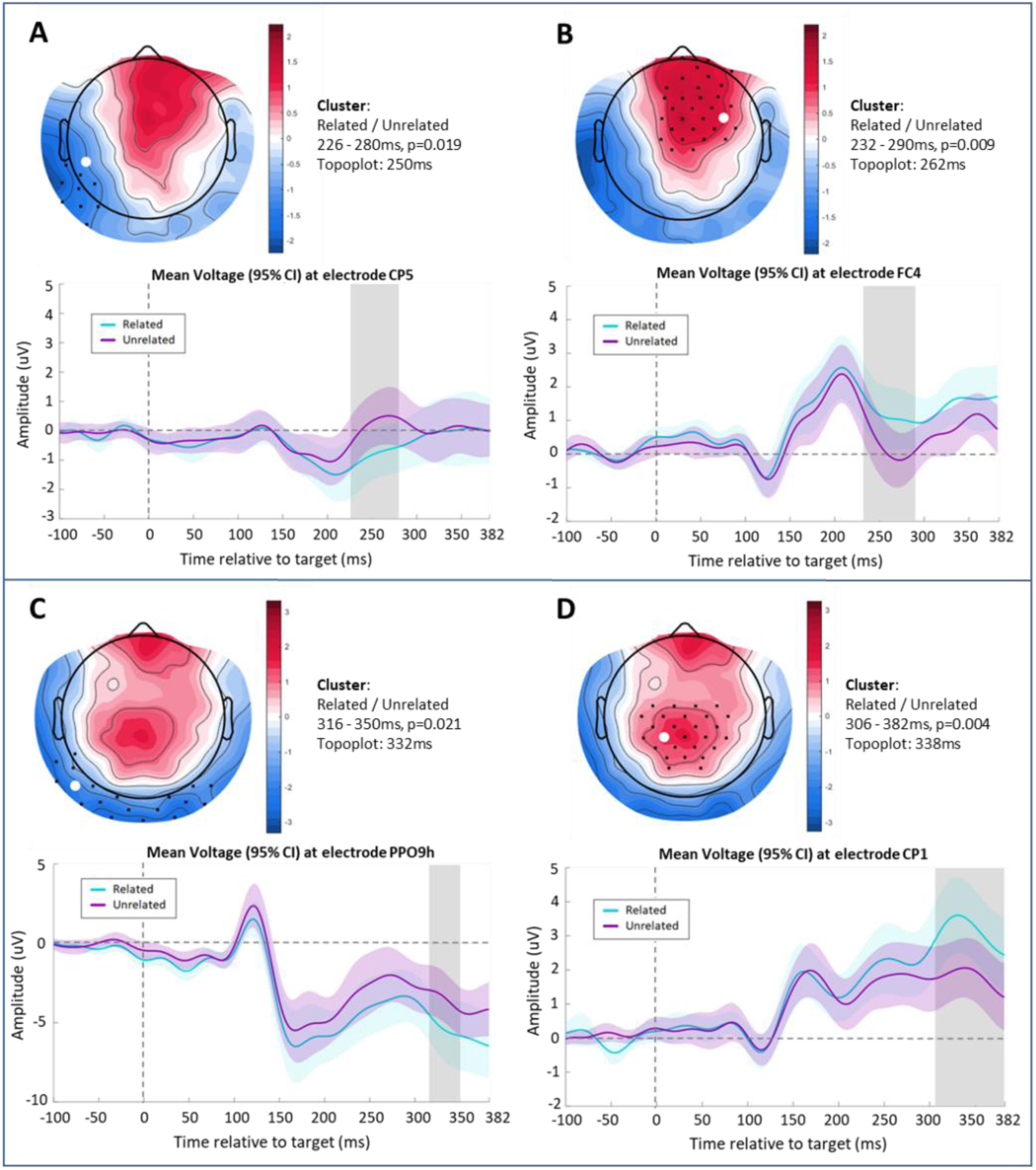
Four main effects from the cluster-based permutation analyses, which contrasted the voltage difference between related and unrelated word-pairs from 0-382ms post-stimulus. ERP scalp topographies revealed two dipolar effects; first, an early fronto-temporal effect at approximately 250ms (A and B); then, a later parieto-occipital effect at around 340ms (C and D). ERP plots show data (mean and shaded 95% confidence interval) from the electrode where the effect was maximal, with the cluster period highlighted in grey.

As an exploratory analysis, and to increase power to detect a potential interaction effect, we tested for the interaction within each of the main effect clusters by averaging per condition and participant across all channels and time points within each main effect cluster. With this approach, the later negative cluster (C in Figure 3) showed a significant interaction (F (1, 21) = 6.679, p = 0.017, ηp^2^= 0.241), reflecting a larger voltage difference between the related and unrelated targets in a high validity context with respect to a low validity context (other clusters p = 0.396; 0.110; 0.273). Bayesian equivalent analyses considered this to be anecdotal evidence for the alternative hypothesis (BF_inclusion_= 1.505), see Figure 4.

**Figure 4:**
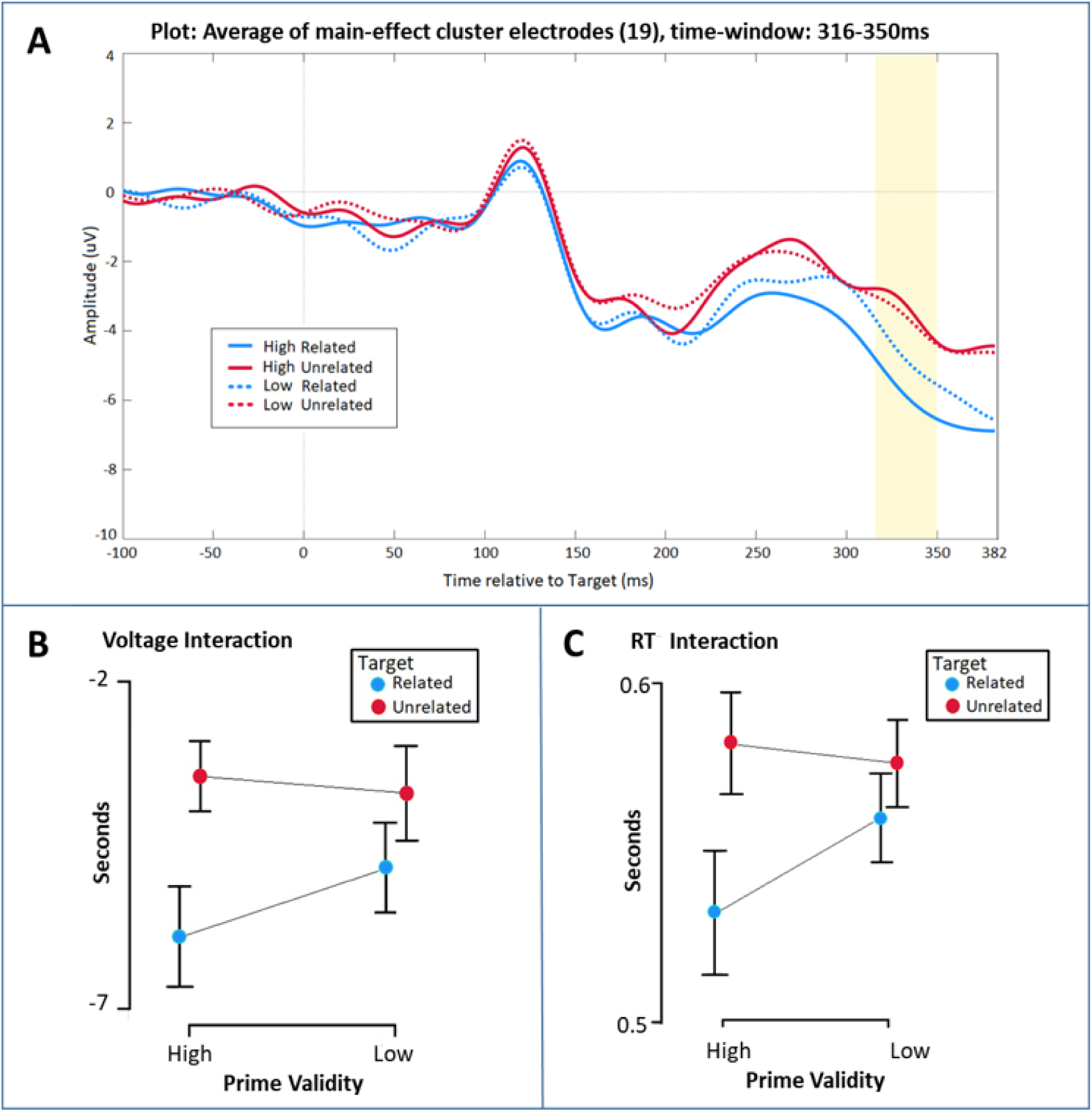
Exploratory analysis to test for the interaction between the four conditions ((HR – HU) – (LR – LU)). The ERP plot in panel A shows the mean of electrodes (19 electrodes) within the 316-350ms cluster found in the main effect analysis (Figure 3, C). Panel B shows the mean for each condition within the same time-window that was analysed with repeated measures ANOVA showing a significant voltage interaction (p = 0.017) with a larger difference in voltage between related and unrelated items in high validity context than low validity context. Panel C shows the significant RT interaction (p = 0.007) presented in Table 2. In this experiment participant’s behaviour (RT; Panel C) showed the same pattern as their ERP responses (Panel B).

#### Source Estimate Analyses

Our pre-registered analyses included whole-brain interaction and main effect contrasts within the time-windows of significant clusters at the sensor level. However, this approach returned no significant clusters at the source level (interaction smallest cluster p = 0.147; main effect smallest cluster p=.067). Furthermore, our preregistered source analyses included regions of interest from the following AAL regions: Left inferior frontal gyrus (LIFG); Left middle temporal gyrus (LMTG); Left superior temporal gyrus (LSTG). However, none of these regions exhibited significant interaction effects or main effects (all FDR corrected p-values > 0.05).

Consequently, for a qualitative visualisation of the source estimates, here we plot the whole-brain thresholded t-values (p<.05) of the source estimate contrasts, uncorrected for multiple comparisons. Specifically, we plot these t-values for the early main effect (Figure 3A&B) and the late main effect (Figure 3C&D) in time windows selected to be entirely within the significant dipolar sensor level clusters (early: 232-280ms; late: 316-350), see Figure 5. The thresholded t-values showed the peak of activity at the Right Middle and Superior Fontal Gyri for the early effect; and the activity peak at the Right Supplementary Motor Area for the late effect, as shown in Figure 5.

**Figure 5:**
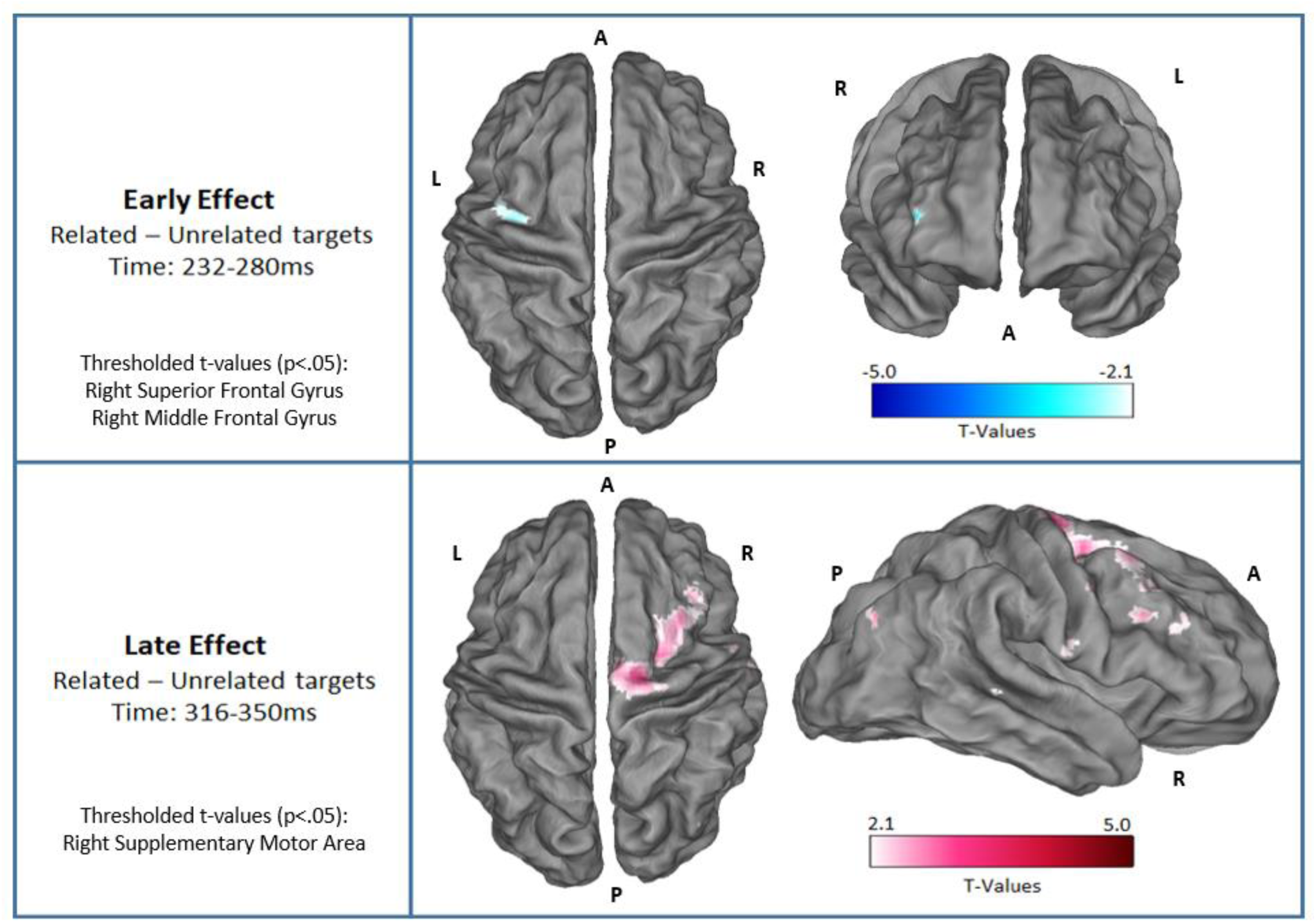
Thresholded t-values (p<.05) of the ERP source estimates over two distinct time windows that corresponded to the early and late ERP effects reported above in Figure 3. In the Figure, the upper panel shows the difference between related and unrelated targets in the early time window (232-280ms), and the lower panel indicates the same difference in a later time window (316-350ms) (thresholded t-values, p < 0.025).

## Discussion

Predictive coding theory posits that the brain generates expectations about upcoming stimuli at varying levels of complexity – from low-level expectations about stimulus properties through to higher-level conceptual expectations. Here, we investigated the behavioural and electrophysiological correlates of such expectations and their violations at two levels of a semantic expectation hierarchy (local and global). First, on the behavioural level, participants of two separate experiments showed evidence of speeded reaction times in related trials relative to unrelated trials, consistent with a local expectation generated about target word identity on the basis of the prime identity. Furthermore, participants generated a more conceptually complex expectation based on the global context (i.e. prime validity) to exhibit greater behavioural facilitation in the high prime validity context than the low prime validity context (Boudewyn, Long, and Swaab, 2015). Importantly, only those individuals who reported conscious strategic expectation showed evidence of behavioural facilitation given by the global context, while those individuals who did not report a conscious strategy only exhibited facilitation as a result of the local context. Together, these behavioural data are consistent with a dissociation between a local expectation about the identity of the target generated by the prime, and a global expectation about the relatedness of the target that necessitates reportable, effortful, and strategic application of expectation. Moreover, the present data provides evidence for a successful replication of the behavioural effect elicited by the same paradigm as implemented by Hutchison (2007), who also found that the magnitude of the global facilitatory effect was modulated by the level of attentional control (i.e. weaker effect in individuals with lower attentional control; Hutchison, 2007). Similarly, our results suggested that only individuals that reported applying an effortful conscious strategy showed the global context effect as mentioned above.

Consistent with this two-stage expectation profile, the ERPs in response to the target words also exhibited a two-stage profile, with an early effect modulated by local expectation (around 250ms) and a later effect modulated by global expectation (around 350ms). These results are broadly consistent with the two-stage profile observed in the auditory oddball local – global paradigm (Bekinschtein et al., 2009), which includes an MMN in an early stage reflecting errors of the local context of the stimuli and a P3b response to errors of the global context given by blocks across the task.

Furthermore, the early effect in the present experiment showed more extreme amplitudes for unexpected targets relative to expected targets, consistent with a prediction error signal, such as the MMN to unexpected/deviant items observed across levels of stimulus awareness (Chennu et al., 2013; Bekinschtein et al., 2009; Faugueras et al., 2012; El Karoui et al. 2014). Moreover, the scalp topography of the early effect has a fronto-central peak, which is consistent with the MMN (Chennu et al., 2013; Bekinschtein et al., 2009; Faugueras et al., 2012), although, its latency is a little longer than seen in some of these previous papers. Additionally, in our source estimation analyses, the early effect was localised to the middle frontal gyrus (Figure 5), whereas in another study the local MMN effect was localised to the temporal parietal junction and prefrontal cortex (Chennu et al., 2013), indicating not entirely overlapping neurocognitive processes. Nevertheless, as we observed behavioural semantic priming (as tracked by the early effect) even for participants who were not making strategic expectations, and due to the shared common features with the MMN (i.e. more extreme for errors and with a fronto-central focus), we consider the early effect to be consistent with an error of local expectation – i.e. expectation based on the identity of the prime, rather than the prime validity. Indeed, the MMN is elicited even by individuals who are not actively attending to the stimuli (Bekinschtein et al., 2009).

The late effect, however, was the opposite of what would be expected for a prediction error signal – i.e. its amplitude was more extreme for expected targets compared to unexpected targets. This *apredictive* pattern is not readily explained by prediction error accounts without appeal to precision-weighting, in which a prediction error is weighted by the system’s confidence in the signal (Chennu et al., 2013; Friston, 2005). Under precision-weighting, all possible patterns of prediction error signals on the scalp are possible, including apredictive patterns as we observed here, as precision may vary freely across task conditions (Kok et al., 2011). For example, Barascud et al. (2016) reported a larger MEG signal for auditory stimuli that become predictable, relative to stimuli that are entirely unpredictable – i.e. an apredictive pattern – that they linked to up-weighting of the expected stimuli by precision (Heilbron, Chait, 2018). Within predictive coding, attention is one specific mechanism that is thought to increase precision (Hohwy, 2012). Therefore, under a predictive coding framework, one can appeal to varying levels of attention across task conditions. Therefore, we could post-hoc theorise that our late apredictive effect reflects individuals paying greater attention to the high validity trials as they have a high level of predictability and paying greater attention to related targets than unrelated targets, as the former fulfil their expectations. Therefore, the relative levels of attention across conditions could interact to generate this apredictive effect. Indeed, consistent with this, 59% of our participants (13/22) self-reported that their strategy was to generate an expectation in the high validity condition only (i.e. “I was trying to guess next word if previous was blue”; where blue was high validity condition).

An alternative interpretation stems from evaluation of our behavioural data. When comparing the behavioural reaction time interaction with the ERP voltage interaction (Table 2 and Figure 4, respectively), both show the same pattern: namely, that the interaction is driven by expected items in a high validity context, showing more extreme values with respect to the other three conditions. This similarity in behaviour and ERP effects suggest that our late ‘error’ effect may simply reflect processing in service of behaviour, whereby sensory signals are routed to goal-driven analogous motor behaviour (Zylberberg et al., 2010). Our late apredictive ERP pattern may therefore not reflect a precision-weighted global prediction error, but more simply the result of the brain routing the incoming information into appropriate behaviour. Under this interpretation, our results are therefore also consistent with interpretations of early ERPs as reflections of prediction error and later ERPs as processes related to conscious access and in support of task demands (e.g. Dehaene & Christen, 2011; Rohaut et al., 2015).

It is possible that other later error signals were also evident in the neural response during our task, including those traditionally linked to the N400 (i.e. peaking approximately 400ms post-target). However, we limited our analyses to the 0 to 382ms time-window post-target so as to avoid muscle artefact created by the pronunciation responses. We chose to use a pronunciation task as our aim was to observe the behavioural effect produced by the manipulation of both the local (relatedness) and global context (prime validity) as implemented by Hutchison (2007). Nevertheless, tasks that don’t produce large muscular artefacts, such as a lexical decision task (LDT) in which individuals only produce motor responses on filler trials, would allow for analysis of the N400 time-window. However, as argued by Hutchison (2007), participants can complete an LDT with a semantic-matching strategy, meaning that after seeing the target they can verify whether it is related to the prime, which could bias their responses as only words can be related and non-words would be, by their nature, unrelated (Hutchison, 2007). Additionally, as we provided a global context by manipulating the proportion of related items across the task, individuals could bias their responses using the validity cue (Keefe & Neely, 1990); for example, primes that were presented in blue (high validity context) were more likely to be related (80%). Therefore, when seeing a blue prime, individuals could judge their response (word/non-word) solely based on the prime, in this case a ‘word’ as most of the word-pairs are related. Instead, using a pronunciation task allows for a purer measure of expectation, with the caveat of limiting the time-window of artefact-free EEG for analysis.

A recent prediction error view on language-related ERPs proposes that the N400 has similar properties to the MMN, as they both are modulated by the predictability of stimuli (i.e. increased ERP amplitude as a prediction-error response) but that their relative latencies indicate prediction-error processing at different levels of stimulus complexity (Bornkessel-Schlesewsky & Schlesewsky, 2019). In our findings, both consecutive effects could be similarly interpreted as reflecting different levels of complexity of precision-weighted prediction error processing across a semantic hierarchy. However, as noted above, appeal to precision-weighting problematically allows for post-hoc explanations of all possible ERP patterns (Bowman, Filetti & Olivers, 2013).

Regarding the source estimation analyses, the early effect was localised to the middle frontal gyrus, which has been previously associated with semantic categorization when compared with passive listening (Noesselt, Shah, Jäncke, 2003). Furthermore, the ERP source estimation analysis for the late effect was localised to the supplementary motor area, consistent with the above interpretation that the late interaction reflects goal-driven routing toward action. Indeed, this area has been linked to speech motor control, verbal working memory, and predictive top-down mechanisms in speech perception (Hertrich, Dietrich, & Ackermann, 2016). However, neither of these two regions were part of our pre-registered hypotheses. Therefore, these source estimates should be interpreted with caution, and future studies with this paradigm will wish to replicate these sources.

In our pre-registered analyses, we also hypothesised that we would observe electrophysiological markers of differential expectations generated by the high and low validity primes, prior to the onset of the target. Specifically, we expected these differential expectations to be reflected in the ERPs, including the slope of a putative slow wave (Chennu et al., 2013), and/or in the power of the EEG in the alpha/beta bands, as these have been previously associated with the precision of expectations (Bauer, Stenner, Friston & Dolan, 2014). However, we found no evidence of any differences in these measures between high and low validity primes prior to target onset. One interpretation is that our specific measures were simply not sensitive enough to detect the differential expectations in these conditions. Indeed, we powered our study to detect the post-target behavioural effect specifically. An alternative interpretation is that expectations were, in fact, not different between the two conditions. Indeed, under predictive coding, the brain is considered to optimize the difference between its expectations and sensory input by updating its internal model (Friston, 2010); hence, it is possible that the optimal means of minimising prediction error in this task is to always predict the related target, regardless of the prime validity. For example, even if one were to consciously expect that an upcoming target will be unrelated (as in a low validity trial), it is simply not possible to accurately predict the identity of that target, as the range of possible unrelated target words is considerable. Therefore, even though predicting the identity of a specific related target had only a ∼22% probability of being correct in a low validity context, it was still more likely than predicting any one of the vast arrays of potential unrelated target words. Future inspection of participants’ meta-cognition in relation to their specific expectations following prime presentation will help speak to this interpretation.

## Conclusions

In conclusion, we here reported ERP evidence of hierarchical matching of semantic expectations to incoming speech. Lower lever expectations based on the local context (i.e. the prime identity) elicited an early and predictive pattern that matches with prediction error accounts. Higher level expectations generated from the global context required awareness of the global rule and the use of a reportable strategy, and were associated with an apredictive pattern that can be interpreted within a precision-weighted prediction error account, or may reflect the routing of sensory signals and their expectations into task-directed behaviour.

## Conflict of Interest

Authors report no conflict of interest.

## Notes

Funding sources: This study was supported by a PhD scholarship given by the Chilean National Agency for Research and Development (Agencia Nacional de Investigación y Desarrollo, ANID), from the Government of Chile; the Medical Research Council (Damian Cruse PI: MR/P013228/1); and the School of Psychology from the University of Birmingham, Birmingham, UK.

https://osf.io/v35te/

### Competing Interest Statement

The authors have declared no competing interest.

